# Pseudotime graph diffusion for post hoc visualization of inferred single-cell trajectories

**DOI:** 10.64898/2026.01.27.702126

**Authors:** Brandon Lukas, Jingbo Pang, Timothy J. Koh, Yang Dai

## Abstract

Visual representations are widely used to interpret trajectories in single-cell data; however, they do not always faithfully capture inferred trajectory structure. As a result, interpretation of cellular dynamics and downstream analyses may be compromised. Here, we present Pseudotime Graph Diffusion (PGD), a lightweight and interpretable post hoc framework for smoothing cell-level features along pseudotime. PGD operates by performing random-walk diffusion on a pseudotime graph, propagating information along inferred trajectory paths to enhance continuity and structure. We demonstrate that PGD-smoothed embeddings improve visualization of increasingly complex inferred trajectories of monocytes and macrophages during wound healing. We further show that PGD extends naturally to trajectory-aware gene expression smoothing and scales to atlas-sized datasets. By improving agreement between visual representations and inferred trajectories, PGD enables more faithful interpretation of dynamic cellular processes.

## 1 Introduction

Single-cell RNA sequencing (scRNA-seq) has enabled the study of dynamic cellular processes at single-cell resolution, providing insights into development, differentiation, and disease-associated state transitions. A central goal of single-cell analysis is to reconstruct continuous cellular trajectories and to order cells along developmental or differentiation axes, motivating the development of trajectory inference and pseudotime estimation methods [1, 2, 3, 4].

In turn, these methods have prompted widespread use of low-dimensional visual representations to project inferred state transitions and to color cells by pseudotime. These representations play a critical role in data interpretation and hypothesis generation and, in some frameworks, serve as substrates from which quantities are derived [5].

Standard visualization methods such as UMAP often represent trajectories for well-ordered and coherent biological processes, such as embryonic development or pancreatic differentiation [6, 7, 8]. However, for less coherent systems, trajectory structures may be obscured. Moreover, when trajectory inference is performed in a higher-dimensional feature space (e.g., principal component space), low-dimensional representations may fail to reflect the inferred structure due to information loss. In such cases, this mismatch can complicate interpretation and downstream analysis.

Several approaches have been specifically proposed to improve trajectory representation in single-cell data. Methods such as PAGA [9], PHATE [10], and DTNE [11] aim to encode global or diffusion-based structure in low dimensions; however, they are not designed to operate in a post hoc setting where inferred trajectories have already been established. Alternatively, one could theoretically visualize a cell-cell transition matrix using any graph layout algorithm; however, this would abstract away cell-level features that could be of interest.

Here, we introduce Pseudotime Graph Diffusion (PGD), a post hoc framework for enhancing agreement between low-dimensional embeddings and existing inferred trajectories or pseudotime estimates. PGD treats a pseudotime graph as a diffusion substrate and smooths cell-level features, such as embeddings or gene expression, along this graph. By diffusing features in a trajectory-aware manner, PGD emphasizes trajectory structure while preserving aspects of the original features.

Using a murine wound-healing dataset, we show that PGD improves visualization of increasingly complex inferred trajectories and naturally extends to trajectory-aware smoothing of gene expression. We further demonstrate that PGD scales to atlas-sized datasets. Our contributions are summarized as follows:

- We introduce PGD, a lightweight, scalable post hoc framework that smooths cellular features along a pseudotime graph to enhance trajectory structure.
- We demonstrate that PGD improves the agreement between low-dimensional visual representations and inferred trajectories, particularly for complex branching structures.
- We show that PGD naturally extends to gene expression smoothing, reducing noise while preserving branch-specific trends.

## 2 Methods

### 2.1 Data and Preprocessing

To evaluate PGD, we analyzed a publicly available scRNA-seq dataset from the Gene Expression Omnibus (GEO; accession GSE203244), consisting of pooled monocytes and macrophages isolated from murine skin wounds at 3, 6, and 10 days post-injury [12].

Data were preprocessed using scanpy [13]. Count matrices from all timepoints were concatenated, and low-quality cells were filtered using median absolute deviation (MAD) thresholds applied to the number of detected genes, total counts, and mitochondrial read fraction. Counts were then normalized and log-transformed. Principal component analysis (PCA) was performed on highly variable genes, and the resulting PC embeddings were corrected for batch effects using Harmony [14].

To demonstrate scalability, we analyzed the Pijuan-Sala mouse gastrulation atlas [15], comprising 116,312 cells profiled across embryonic stages E6.5–E8.5. The atlas was obtained via the MouseGastrulationData Bioconductor package [16]. We retained the published iterative fastMNN-corrected PCs from the atlas and excluded cells flagged as stripped nuclei or doublets.

### 2.2 Trajectory Inference with Lamian

Trajectory structure and pseudotime were inferred using Lamian, a statistical framework for pseudotemporal trajectory analysis [17]. Lamian constructs a cluster-based trajectory graph in a low-dimensional feature space and assigns pseudotime values along inferred paths.

As input, we provided Harmony-corrected PC embeddings along with cell-level metadata indicating sampling timepoints. The Lamian cluster with the largest proportion of cells from day 3 post-injury was specified as the pseudotime origin. To model varying branching complexities, Lamian was run across a range of cluster resolutions (5–10 clusters), and the resulting trajectory graphs and pseudotime assignments were used as downstream inputs for PGD.

For the gastrulation atlas, the same Lamian pipeline was applied to the published fastMNN-corrected PCs, with embryonic stage E6.5 specified as the pseudotime origin. We report scalability results at a cluster resolution of 17.

### 2.3 Pseudotime Graph Diffusion

Fig. 1 provides an overview of PGD. Let *C* = (*c*_1_, …, *c*_*n*_) denote an ordering of *n* cells by inferred pseudotime, and let *X* ∈ℝ ^*n*×*d*^ denote the corresponding cell-level feature matrix with *d* features per cell. PGD constructs an undirected cell graph by connecting cells that are nearby along the pseudotime ordering. Specifically, the edge set is defined using a sliding window as

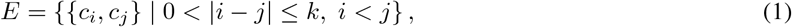

where *k* ≥1 is the window radius. For branched trajectories, this construction is applied separately within each branch-specific ordering, and the resulting edge sets are combined by union.

**Figure 1.**
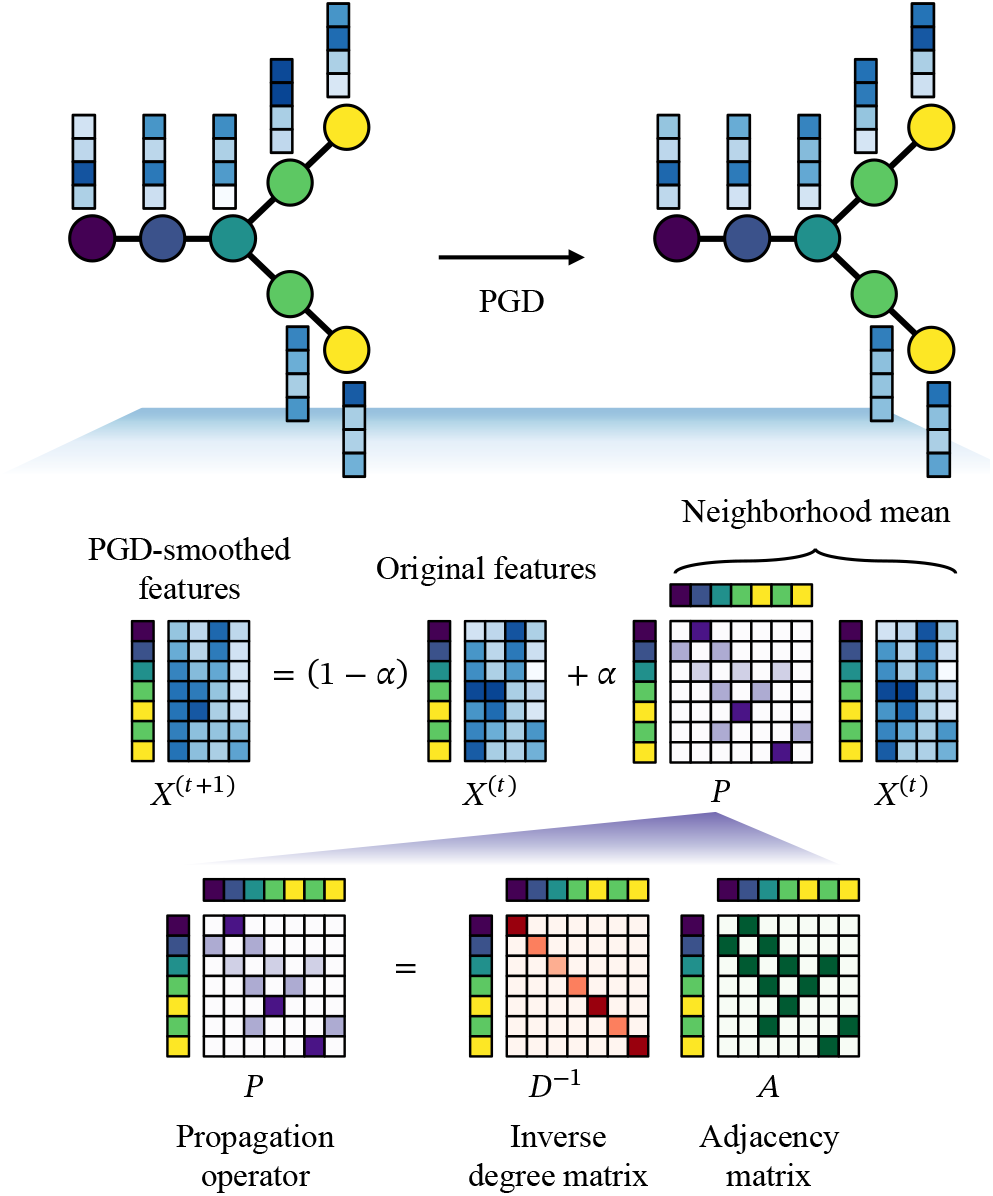
Schematic of PGD.

PGD updates cell features using a lazy random-walk diffusion on this graph. At diffusion step *t* = 0, 1, 2, …, features are updated according to

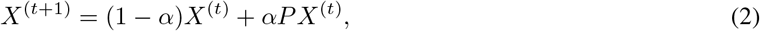

where *X*^(*t*)^ denotes the feature matrix at step *t, A* ∈ ℝ ^*n*×*n*^ is the symmetric adjacency matrix of the graph, *D* is the diagonal degree matrix with entries *D*_*ii*_ = Σ _*j*_ *A*_*ij*_, and

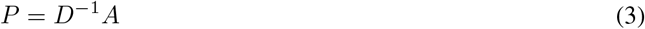

is the random-walk propagation operator. The parameter *α* ∈[0, 1] controls the diffusion strength by weighting contributions from neighboring cells relative to the original features. When *A* has no self-loops, *PX*^(*t*)^ corresponds to the local neighborhood mean of each cell’s features.

Defining the combined propagation operator

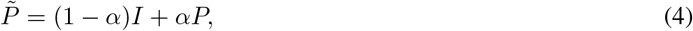

the diffusion process can be written in matrix-power form as

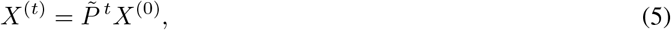

where the exponent *t* denotes the number of diffusion steps (matrix power), not transposition. Unless otherwise stated, we use *k* = 50, *α* = 0.5, and a single diffusion step (*t* = 1).

### 2.4 Comparison with MAGIC

We compared PGD against MAGIC [18], a widely used diffusion-based smoothing and imputation method for scRNA-seq data. MAGIC performs diffusion over an expression-derived affinity graph, whereas PGD diffuses over a pseudotime graph. We applied MAGIC using its default parameters. For a controlled comparison, we configured PGD with the same number of diffusion steps (*t* = 3) and set the pseudotime window radius to *k* = 8. Because PGD connects each cell to neighbors in both directions along pseudotime (up to 2*k* neighbors), this setting yields a maximum neighborhood size that is comparable to the default maximum of 15 neighbors in MAGIC.

### 2.5 Benchmarking

To assess scalability, we benchmarked PGD on both the gastrulation atlas and the wound-healing dataset. Wall time and peak resident set size (RSS) were measured for pseudotime graph construction and feature diffusion. Benchmarks were run on a single MacBook (Apple M-series, 8 physical cores, 17.2 GB RAM; Python 3.12, NumPy 2.4, SciPy 1.17).

## 3 Results

### 3.1 Monocyte and Macrophage Wound-Healing Dataset

After preprocessing, the wound-healing dataset consisted of 2,848 monocytes and macrophages distributed across days 3 (34.4%), 6 (19.6%), and 10 (45.9%) post-injury (Fig. 2). When visualized using UMAP, the Harmony-corrected PC embeddings display a continuous spectrum of cell states without distinct clustering. While cells from different timepoints occupy generally distinct regions, suggestive of temporal progression, the standard UMAP layout lacks a clearly defined trajectory structure. This lack of visual coherence contrasts with more structured developmental systems, making this dataset an excellent test case for evaluating whether PGD can enhance the representation of trajectories inferred in higher-dimensional spaces.

**Figure 2.**
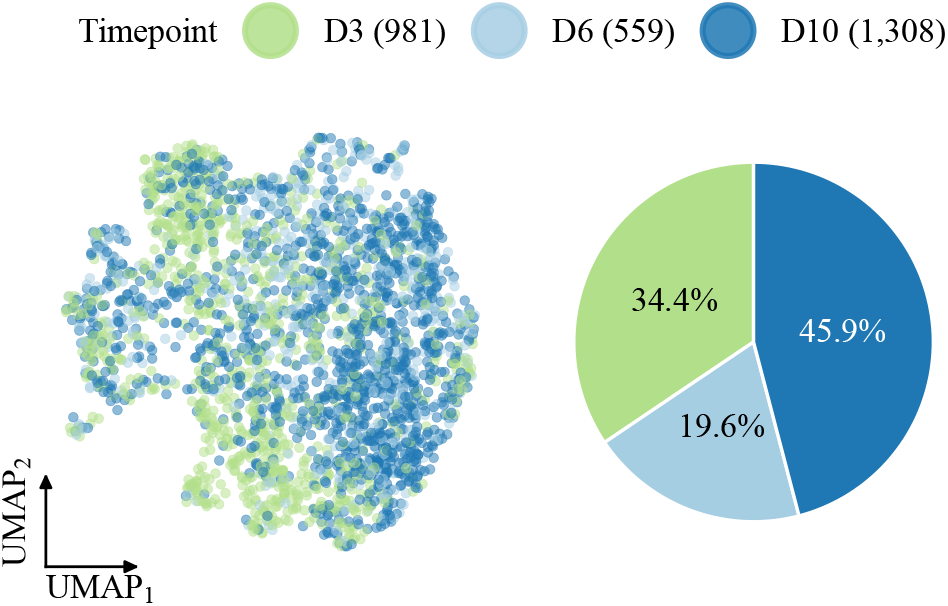
UMAP of the monocyte/macrophage wound-healing dataset colored by timepoint (left) and timepoint composition shown as a pie chart (right).

### 3.2 PGD-Smoothed Embeddings Improve Visualization of Inferred Trajectories

We first evaluated whether PGD improves visual representations of inferred trajectories. Since UMAP is a standard approach for visualizing single-cell data, we applied PGD to Harmony-corrected PC embeddings using Lamian-inferred trajectory graphs at varying cluster resolutions, then compared UMAP layouts generated before and after smoothing (Fig. 3).

**Figure 3.**
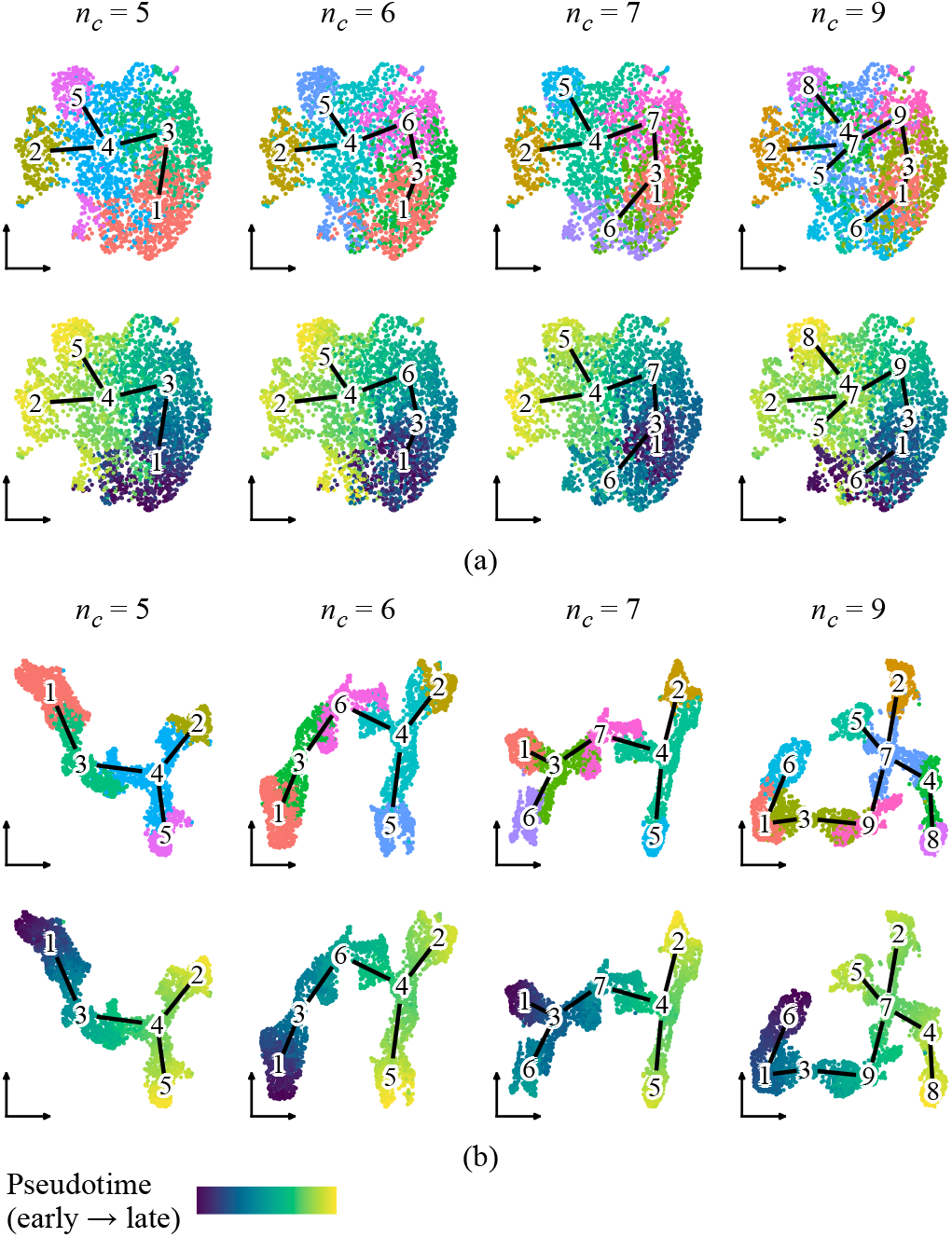
UMAP layouts generated from (a) original embeddings and (b) PGD-smoothed embeddings. Cells are colored by Lamian cluster or pseudotime. Numbers mark Lamian cluster centroids. Edges indicate the Lamian-inferred trajectory graph.

Lamian inferred progressively more complex trajectory structures using five, six, seven, and nine clusters; however, the UMAP layout constructed from embeddings before PGD remained fixed across these settings. When cells were colored by Lamian cluster or pseudotime, this static layout did not clearly reflect the inferred trajectories, particularly at higher cluster resolutions (Fig. 3a). As a result, cells distant along pseudotime sometimes appeared proximal in the embedding, forcing a mismatch between the visual representation and the inferred dynamics.

In contrast, UMAP layouts constructed from embeddings after diffusion were specific to each inferred trajectory, exhibiting increased continuity and organization along inferred paths. When cells were colored by pseudotime, the layout displayed smoother and more coherent gradients, and separation between trajectory branches became visually apparent (Fig. 3b). Notably, in the nine-cluster trajectory where clusters 5 and 6 originally overlapped despite lying at opposite ends of the pseudotime continuum, PGD clearly separated them while keeping them within the same global neighborhood.

These results suggest that PGD preserves coarse-scale relationships from the original embeddings while enhancing trajectory-consistent structure. By diffusing low-dimensional embeddings along a pseudotime graph, PGD improves the agreement between visual representations and inferred trajectory structures, enabling more faithful interpretation of dynamic processes.

### 3.3 Effect of Diffusion Strength Parameter

We next examined how the diffusion strength parameter *α* influences the visual structure of smoothed embeddings. Using the Lamian-inferred trajectory with five clusters, we applied PGD to Harmony-corrected PC embeddings while incrementally increasing *α* from 0.05 to 0.80 and visualized the results using UMAP (Fig. 4).

**Figure 4.**
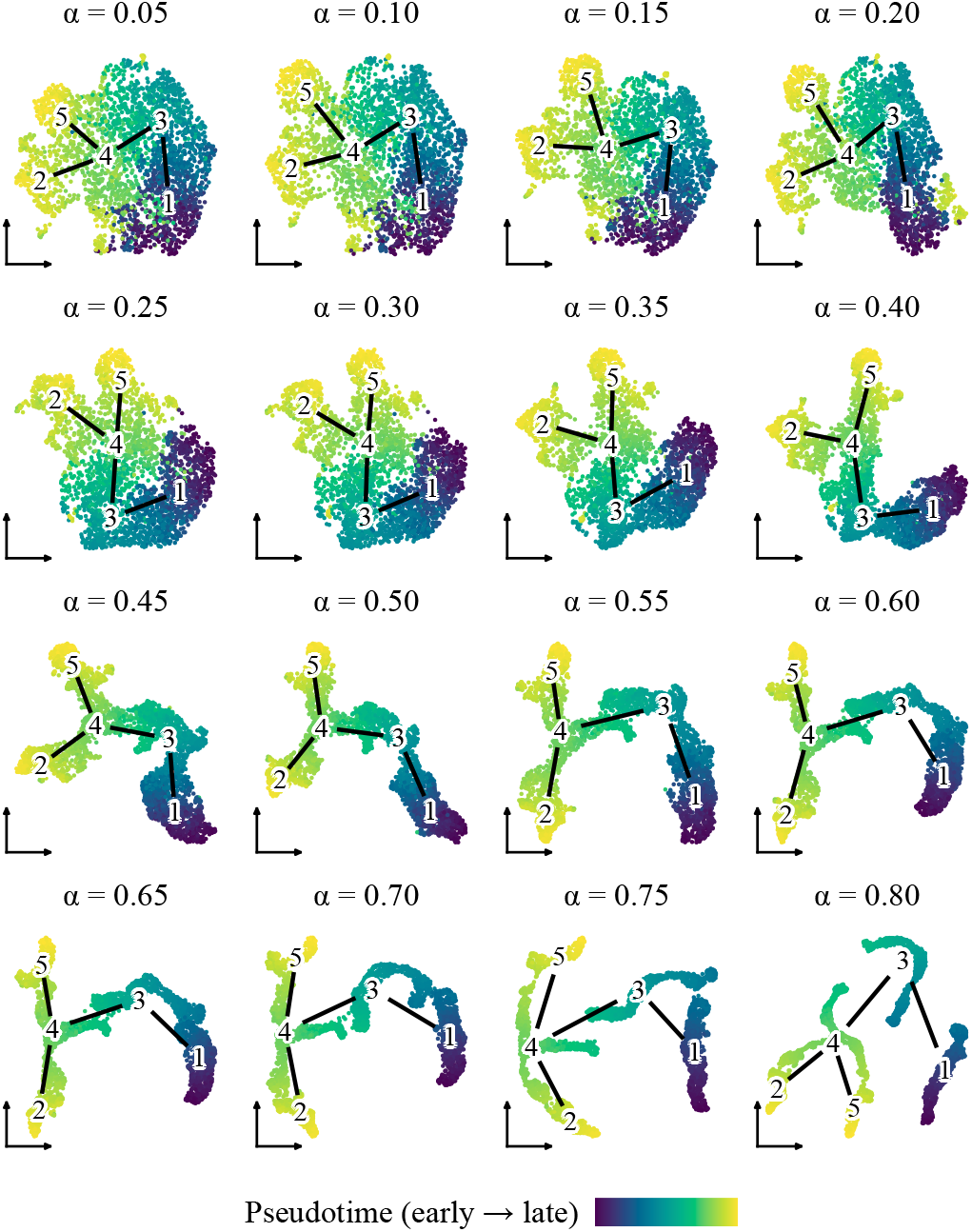
UMAP layouts generated from PGD-smoothed embeddings for incrementally increasing *α*. Cells are colored by pseudotime. Numbers mark Lamian cluster centroids. Edges indicate the Lamian-inferred trajectory graph.

The visualization response varied distinctly across three regimes. At low diffusion strengths (*α <* 0.30), smoothing was subtle, and embeddings retained a geometry largely similar to the original input. At moderate strengths (*α* ≈ 0.30–0.50), trajectory structure became visually dominant; pseudotime gradients appeared smoother, and branches became more coherent. Conversely, at high strengths (*α >* 0.50), the embeddings began to collapse into one-dimensional artifacts, abstracting away local cell-level variation in favor of the global trajectory backbone.

These results confirm that *α* offers an interpretable mechanism to tune the balance between data fidelity and trajectory emphasis. For most visualization tasks, intermediate values provide an effective compromise that enhances structural clarity without forcing over-simplification.

### 3.4 PGD Extends to Trajectory-Aware Smoothing of Gene Expression

We next examined whether PGD can extend to smoothing gene expression along inferred trajectories. This is a useful task in single-cell analysis because scRNA-seq measurements are inherently sparse and noisy due to dropout and technical variability, which can obscure fine-scale expression dynamics.

To evaluate this, we focused on genes known to drive monocyte and macrophage dynamics during wound healing: *Il1b, Nfkbia, Arg1*, and *H2-Ab1*. In early wounds (0–3 days post-injury), monocytes and macrophages exhibit a pro-inflammatory phenotype characterized by high expression of *Il1b* and *Nfkbia* [19, 20, 12]. As wounds transition to the proliferative phase (3–7 days), cells shift toward a repair phenotype, upregulating genes such as *Arg1* [21, 19, 12, 22]. In parallel, expression of the MHC class II gene *H2-Ab1* increases and reflects antigen presentation functions [12].

Using the Lamian-inferred trajectory with nine clusters, we applied PGD to smooth the expression of these markers along pseudotime. We then visualized the raw (normalized), PGD-smoothed, and MAGIC-smoothed expression patterns on UMAP and as a function of pseudotime (Fig. 5).

**Figure 5.**
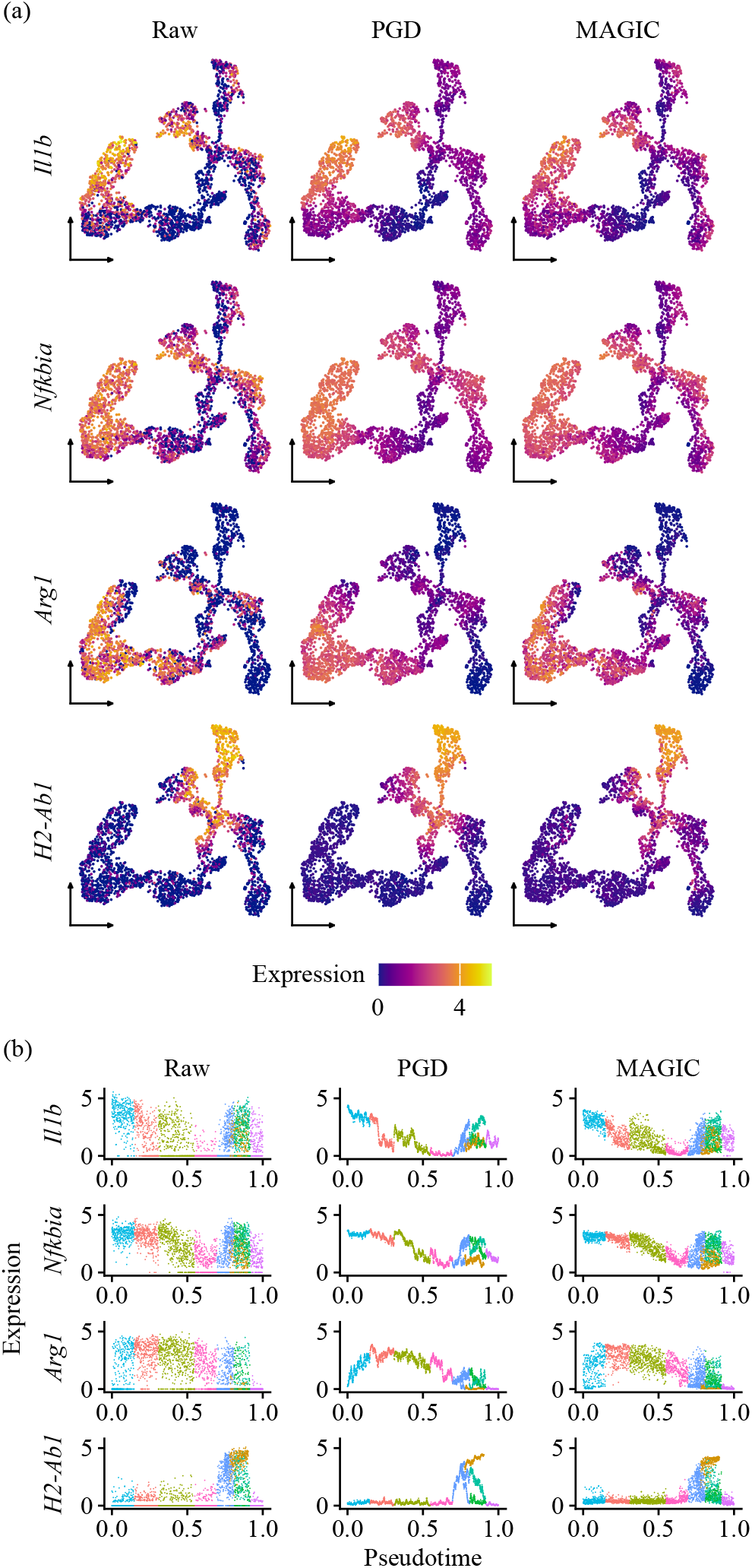
Log-normalized expression of representative genes (a) on UMAP and (b) as a function of pseudotime. In (b), cells are colored by Lamian cluster.

Both PGD and MAGIC effectively reduced high-frequency noise from the raw data, yielding highly correlated results overall (Pearson correlation *ρ* = 0.882 across matched cell-gene pairs for the four genes). However, subtle spatial differences emerged on UMAP. For instance, *H2-Ab1* expression in PGD-smoothed data appeared more tightly confined to the top trajectory branch, whereas MAGIC-smoothed expression diffused more broadly across adjacent branches (Fig. 5a). This localization is consistent with previous RNA velocity analyses identifying *H2-Ab1* as a driver for antigen-presenting clusters [12].

Differences became more pronounced when visualizing expression as a function of pseudotime (Fig. 5b). PGD-smoothed expression displayed sharper branch-specific separation and cleaner pseudotemporal gradients. In contrast, MAGIC-smoothed expression appeared noisier along the pseudotime axis and more blended between branches. Notably, for *Nfkbia*, PGD clearly resolved two distinct expression levels across separate branches in late pseudotime (approximately 0.7–0.9), a separation that was less distinct in the MAGIC-smoothed result. This separation aligns with the identification of *Nfkbia* as a driver gene along early-to late-stage clusters [12].

Together, these results illustrate that PGD provides a trajectory-aware approach to gene expression smoothing. By enforcing continuity along the inferred pseudotime graph, PGD emphasizes branch-specific dynamics over global similarity, facilitating clearer interpretation of gene expression patterns along cellular trajectories.

### 3.5 PGD Scales to Atlas-Sized Datasets

To assess scalability, we benchmarked PGD on the gastrulation atlas (116,312 cells) alongside the wound-healing dataset (2,848 cells) for reference. PGD completed end-to-end on the 116k-cell atlas in under two seconds with peak memory below 2 GB. Between this and the wound-healing benchmark, runtime grew approximately proportionally with cell count while memory grew sublinearly (Table 1).

**Table 1.** PGD runtime and peak memory usage.

| Step | Wound-healing<br>(2,848 cells) | Gastrulation<br>(116,312 cells) |
| --- | --- | --- |
| Graph construction | 0.02 s | 0.82 s |
| Feature diffusion | 0.01 s | 0.68 s |
| **Total wall time** | **0.03 s** | **1.50 s** |
| Peak memory (RSS) | 0.50 GB | 1.66 GB |

Qualitatively, the PGD-smoothed embedding more clearly resolves the branching structure of the inferred gastrulation trajectory without disrupting the geometry of the original embedding (Fig. 6; cluster compositions in Table 2). For example, clusters 17 (primitive streak and early derivatives; predominantly E7.0–E7.25) and 16 (surface ectoderm and endoderm; predominantly E8.25–E8.5) detach, and cluster 2 (neuro-mesodermal progenitors and neural derivatives) emerges as a more visually distinct bifurcation point.

**Table 2.** Lamian cluster composition for the mouse gastrulation atlas.

| Cluster | Cells | Top cell types<sup>a</sup> | Peak stages<sup>b</sup> |
| --- | --- | --- | --- |
| 1 | 8,860 | Epiblast (49%); Primitive Streak (35%); Rostral neurectoderm (15%) | E7.0–E7.5 |
| 2 | 7,165 | Forebrain/Midbrain/Hindbrain (28%); Neuro-mesodermal progenitor (26%); Spinal cord (15%) | E8.25–E8.5 |
| 3 | 11,655 | Epiblast (86%); Primitive Streak (13%) | E7.0–E7.25 |
| 4 | 5,062 | Mesenchyme (64%); Allantois (17%) | E8.0–E8.5 |
| 5 | 6,757 | Nascent mesoderm (49%); Mixed mesoderm (17%); Paraxial mesoderm (12%) | E7.0–E7.25 |
| 6 | 9,100 | Forebrain/Midbrain/Hindbrain (29%); Caudal epiblast (21%); Rostral neurectoderm (18%) | E7.75–E8.25 |
| 7 | 11,759 | ExE ectoderm (100%)<sup>c</sup> | E7.0–E7.75 |
| 8 | 8,192 | Haematoendothelial progenitors (27%); Mesenchyme (21%); ExE mesoderm (13%); Allantois (12%); Endothelium (10%) | E7.75–E8.5 |
| 9 | 2,188 | Blood progenitors 2 (64%); Blood progenitors 1 (32%) | E7.5–E8.0 |
| 10 | 4,283 | Erythroid3 (63%); Erythroid2 (25%); Erythroid1 (12%) | E8.25–E8.5 |
| 11 | 9,265 | ExE endoderm (100%) | E7.5–E7.75 |
| 12 | 1,551 | Visceral endoderm (76%); Parietal endoderm (15%) | E7.0–E7.5 |
| 13 | 12,450 | Paraxial mesoderm (22%); Intermediate mesoderm (21%); Pharyngeal mesoderm (21%); Somitic mesoderm (13%); ExE mesoderm (12%) | E8.0–E8.5 |
| 14 | 2,245 | Gut (59%); Notochord (17%); Surface ectoderm (12%) | E7.75–E8.5 |
| 15 | 3,642 | Erythroid1 (67%); Blood progenitors 2 (33%) | E7.75–E8.25 |
| 16 | 4,717 | Surface ectoderm (49%); Definitive endoderm (16%); Gut (12%) | E8.25–E8.5 |
| 17 | 7,421 | Primitive Streak (35%); Rostral neurectoderm (30%); Nascent mesoderm (13%); Anterior Primitive Streak (11%) | E7.0–E7.25 |

<sup>a</sup>Listed in decreasing order; restricted to types comprising at least 10% of the cluster.<sup>b</sup>Stages comprising at least 15% of the cluster.<sup>c</sup>ExE: extraembryonic.

**Figure 6.**
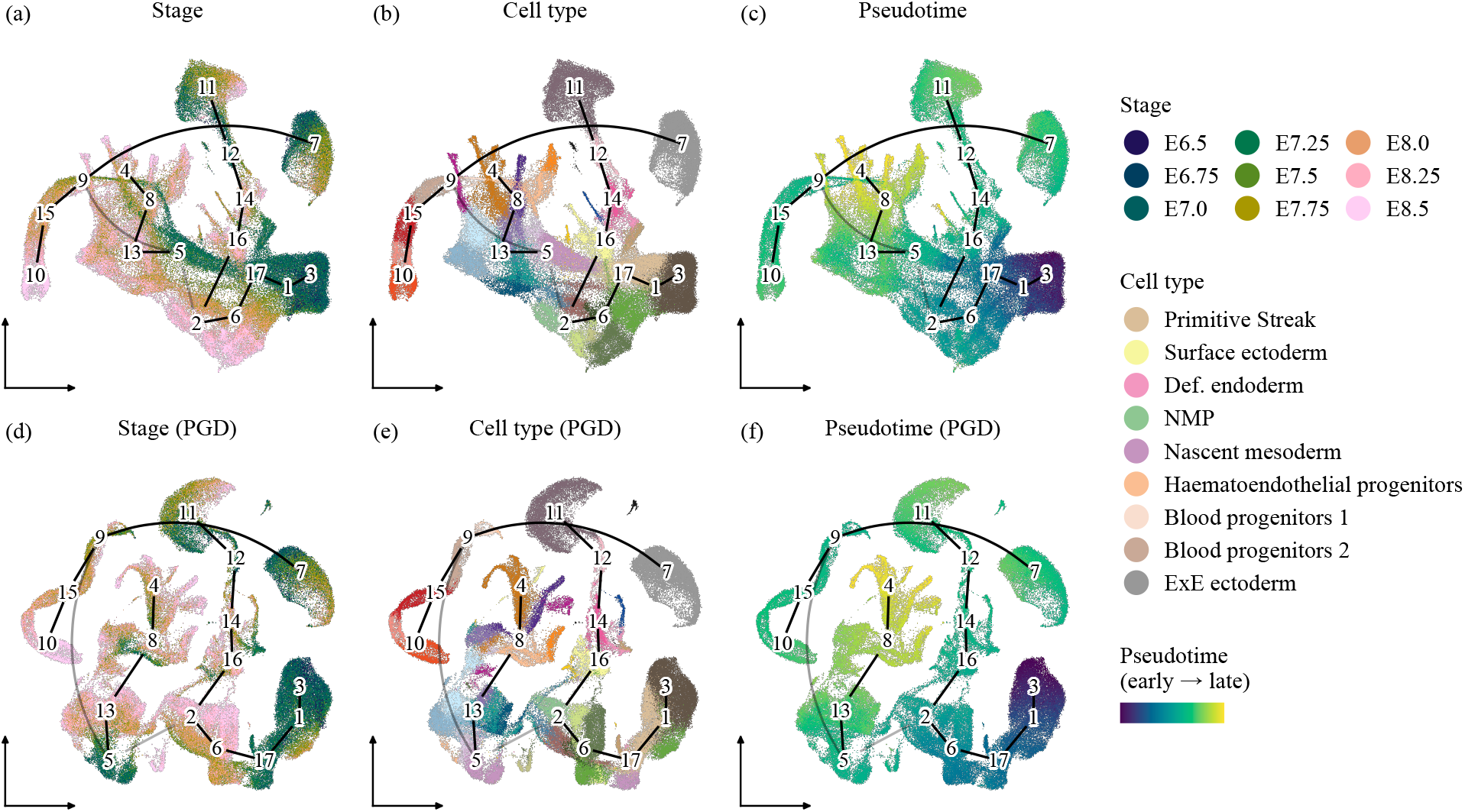
UMAP layouts of the mouse gastrulation atlas (116,312 cells) generated from (a–c) original embeddings and (d–f) PGD-smoothed embeddings. Cells are colored by (a, d) embryonic stage (E6.5–E8.5), (b, e) cell type, and (c, f) pseudotime. Numbers mark Lamian cluster centroids. Edges indicate the Lamian-inferred trajectory graph, with opacity proportional to detection rate across permutations. Cell-type palette from [15]; 9 of 37 cell types are shown in the legend for space. Def., definitive; NMP, neuro-mesodermal progenitor; ExE, extraembryonic.

At the same time, two inferred edges (5–9 and 9–7) connect cluster centroids that remain visually distant in both the original and PGD-smoothed layouts. Edge 5–9 had a low detection rate across 1,000 permutations (0.27), indicating a weakly supported inference, although the connection between early mesoderm (cluster 5) and blood progenitors (cluster 9) is biologically plausible via hematoendothelial progenitors [15]. In contrast, edge 9–7, despite a high detection rate (0.99), connects extraembryonic ectoderm (cluster 7) to blood progenitors, lineages of distinct developmental origin with no expected biological transition. Because trajectory inference methods such as Lamian construct a connected backbone over all clusters, isolated clusters can be reproducibly attached to their nearest neighbor in feature space even when no lineage relationship exists. Rather than warping the embedding to fit such edges, PGD defers to the underlying geometry. Together, these results support the practical applicability of PGD to atlas-scale datasets and illustrate that the deference of PGD to embedding geometry preserves biologically meaningful structure even when the supplied trajectory contains spurious connections.

## 4 Discussion

Here, we demonstrated that PGD provides a simple post hoc strategy for improving the alignment between inferred trajectories and low-dimensional visual representations. By diffusing cell-level features along a pseudotime graph, PGD enhances trajectory-consistent structure and continuity to facilitate interpretation of cellular dynamics. Across increasing levels of trajectory complexity, PGD produced smoother pseudotime gradients and clearer branch separation, reflecting the explicit constraint of diffusion along inferred trajectory paths. We further demonstrated that PGD naturally extends to smoothing gene expression along trajectories, emphasizing pseudotime-dependent and branch-specific expression gradients. Finally, PGD scaled to atlas-sized datasets, supporting its practical applicability to modern scRNA-seq workflows.

PGD operates entirely downstream of trajectory inference and pseudotime estimation methods, refining cell-level features while leaving underlying trajectories and pseudotime unchanged. This modular design ensures PGD can be readily integrated with diverse trajectory inference frameworks without disrupting established upstream workflows.

Several limitations should be noted. First, the evaluations presented here are primarily qualitative. Second, PGD currently diffuses each feature independently, whereas many biological features exhibit structured dependencies that could be leveraged for improved smoothing. Third, we restricted our analysis to unweighted, undirected pseudotime graphs, although PGD can be generalized to weighted or directed graphs and to more general cell-cell transition matrices. Finally, PGD does not infer trajectories or generate embeddings de novo, and its effectiveness likely depends on the quality and consistency of the supplied trajectory and embeddings. More broadly, PGD offers a simple and extensible framework for enhancing the interpretability of single-cell trajectories.

## Acknowledgments

The authors acknowledge the use of AI-assisted tools, including ChatGPT, Gemini, and Claude, for language editing and grammar enhancement; Claude and GitHub Copilot for accelerating code development and visualization; and Claude for research and ideation, including identifying related work and providing feedback on ideas. The authors take full responsibility for the content of the manuscript.

